# Excitatory Cortical Neurons from CDKL5 Deficiency Disorder Patient-Derived Organoids Show Early Hyperexcitability Not Identified in Neurogenin2 Induced Neurons

**DOI:** 10.1101/2024.11.11.622878

**Authors:** Madison R. Glass, Dosh Whye, Nickesha C. Anderson, Delaney Wood, Nina R. Makhortova, Taryn Polanco, Kristina H. Kim, Kathleen E. Donovan, Lorenzo Vaccaro, Ashish Jain, Davide Cacchiarelli, Liang Sun, Heather Olson, Elizabeth D. Buttermore, Mustafa Sahin

## Abstract

CDKL5 deficiency disorder (CDD) is a rare developmental and epileptic encephalopathy resulting from variants in cyclin-dependent kinase-like 5 (CDKL5) that lead to impaired kinase activity or loss of function. CDD is one of the most common genetic etiologies identified in epilepsy cohorts. To study how CDKL5 variants impact human neuronal activity, gene expression and morphology, CDD patient-derived induced pluripotent stem cells and their isogenic controls were differentiated into excitatory neurons using either an NGN2 induction protocol or a guided cortical organoid differentiation. Patient-derived neurons from both differentiation paradigms had decreased phosphorylated EB2, a known molecular target of CDKL5. Induced neurons showed no detectable differences between cases and isogenic controls in network activity using a multielectrode array, or in MAP2+ neurite length, and only two genes were differentially expressed. However, patient-derived neurons from the organoid differentiation showed increased synchrony and weighted mean firing rate on the multielectrode array within the first month of network maturation. CDD patient-derived cortical neurons had lower expression of CDKL5 and HS3ST1, which may change the extracellular matrix around the synapse and contribute to hyperexcitability. Similar to the induced neurons, there were no differences in neurite length across or within patient-control cell lines. Induced neurons have poor cortical specification while the organoid derived neurons expressed cortical markers, suggesting that the changes in neuronal excitability and gene expression are specific to cortical excitatory neurons. Examining molecular mechanisms of early hyperexcitability in cortical neurons is a promising avenue for identification of CDD therapeutics.

**Graphical Abstract:** 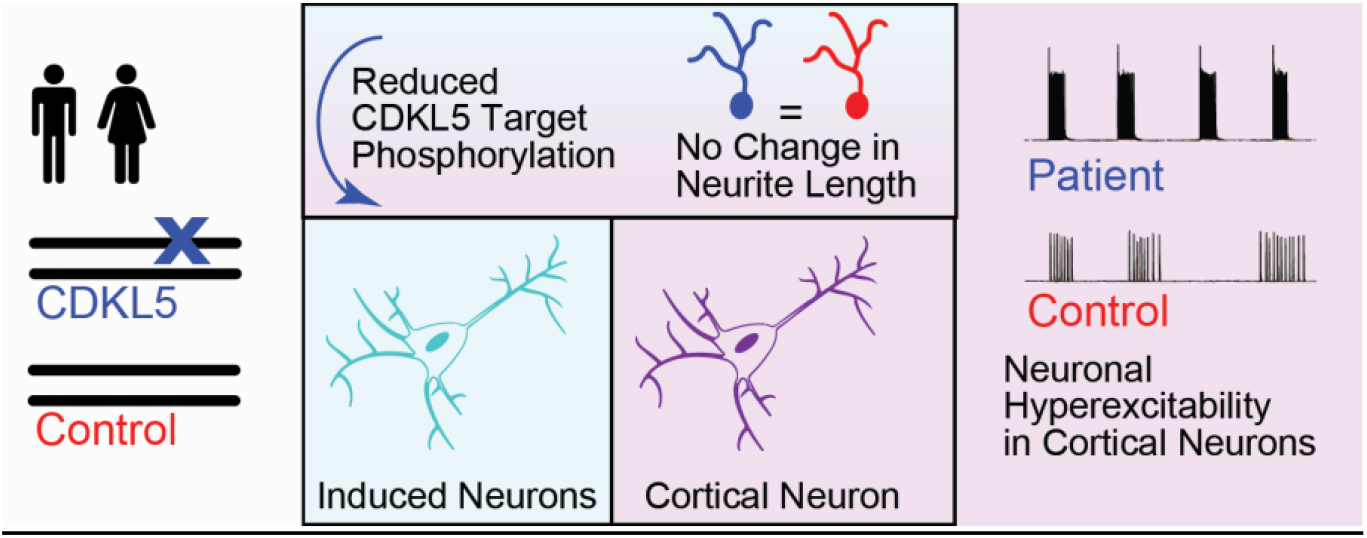

## Introduction

CDKL5 deficiency disorder (CDD) is a rare encephalopathy occurring in 1 in every 40,000 live births (Olson et al. 2019). CDKL5 (cyclin-dependent kinase-like 5) is a serine-threonine kinase located on the X chromosome, and variants disrupting its kinase function result in drug-resistant seizures early in infancy along with neurodevelopmental delay and motor dysfunction (Fehr et al. 2013; Leonard et al. 2022).

The effects of loss of CDKL5 function on neuron development and on the excitatory/inhibitory balance in the brain have been the subject of investigation in model systems ranging from Drosophila to rodents to human induced pluripotent stem cells (iPSCs). To determine if specific neuronal subtypes are more significantly affected by loss of CDKL5 function and thus contribute to network imbalances, groups have used rodent models that target Cdkl5 loss of function in specific cell types. One recent study revealed that loss of *Cdkl5* in adult mouse glutamatergic neurons was sufficient to induce seizures through an increase in BDNF-TrkB signaling (Z.-A. Zhu et al. 2023). Another group revealed that both glutamatergic and GABAergic neurons, but not astrocytes, express Cdkl5 and a target of Cdkl5 kinase activity, EB2 (phosphorylated at Serine222) (Silvestre et al. 2024), suggesting that CDKL5 has cell type specific effects.

Studies in mouse models and human CDD patient-derived iPSCs have also uncovered neuronal mechanisms that may contribute to disease pathology in CDD. These include changes in neurite outgrowth, synapse formation (Y.-C. Zhu et al. 2013; Ricciardi et al. 2012; Y.-C. Zhu and Xiong 2019), neural progenitor proliferation (Fuchs et al. 2022), and transcriptional regulation (Mari et al. 2005; Trazzi et al. 2016; Van Bergen et al. 2022). Though additional targets of CDKL5 may still be uncovered, CDKL5 kinase activity is known to target proteins involved in microtubule assembly, including MAP1S, EB2, and ARHGEF2 (Baltussen et al. 2018; Muñoz et al. 2018), and neuronal function, including voltage-gated calcium channel CaV2.3 (Sampedro-Castañeda et al. 2023). These findings broadly implicate CDKL5 in neuronal function.

To uncover cell autonomous changes in human glutamatergic cortical neurons with loss of CDKL5 function, previous studies used CDD patient-derived iPSCs with variants across several domains of the gene. The CDD patient iPSCs were differentiated into adherent neurons, neural progenitor cells, and human cortical organoids (Negraes et al. 2021). Phenotypic profiling of cortical organoid-derived neurons revealed hyperexcitability in CDD neurons, as well as increased neurite length, increased mTOR activity, and reduced phosphorylation of CDKL5 targets (Wu et al. 2022; Negraes et al. 2021). However, others have shown that CDD patient iPSC-derived cortical neurons have reduced neurite length *in vitro* (Van Bergen et al. 2021), and rodent studies revealed a decrease in mTOR activity with loss of CDKL5 function (Amendola et al. 2014). This left us with questions about whether methods for differentiation or neuronal subtypes contribute to differences in phenotype presentation in each model.

We have previously generated iPSC lines from CDD patients who harbor truncating variants in CDKL5 between amino acids 172 to 781, along with isogenic controls (Chen et al. 2021). We hypothesized that differentiating these iPSC lines into glutamatergic cortical neurons would result in increased neuronal excitability underpinned by changes in synaptic gene expression and neurite length.

To test this hypothesis, we characterized glutamatergic neurons derived via induced Neurogenin2 (NGN2) overexpression (iNs) (Zhang et al. 2013) and neurons derived from a guided cortical organoid differentiation (ACTX) (Whye et al. 2024) to identify potential CDKL5 variant effects that are consistent across protocols. Neurons from both methods of differentiation were used to assess changes in neuronal function, morphology, and gene expression. Unexpectedly, we did not observe any consistent changes in neurite length across patient lines in either differentiation method, but we did identify early hyperexcitability in CDD patient-derived lines when using the ACTX protocol, and not the iN protocol. Using transcriptomic analysis, we found that the ACTX-derived neurons more closely resembled primary cortical neurons than the iNs, suggesting that cortical identity is required to produce CDD-specific changes in neuronal function. Taken together, our study suggests that hyperexcitability of cortical neurons is a consistent cellular phenotype across multiple CDD patients *in vitro* and may be an endpoint that can be utilized for assessing potential therapeutics.

## Results

We have previously generated iPSCs from 6 CDD patients with single nucleotide variants in *CDKL5* (Chen et al. 2021). Male CDD patients express mosaic somatic variants, and isogenic controls were established by screening iPSC clones with no variants during the reprogramming process (Table 1). Female CDD iPSC lines were established by selecting clones expressing either the variant or control *CDKL5* sequence on the active X chromosome, which is regulated by random X inactivation. To develop an *in vitro* model system for assessing cell autonomous effects of loss of CDKL5 function on glutamatergic cortical neurons, we first chose to differentiate two male CDD iPSC lines into cortical neurons using Neurogenin2 overexpression (iNs). The male donors were selected to control for sex effects, to eliminate the need for screening for X-inactivation switching in female lines, and to potentially increase the likelihood of a strong phenotype since males have more severe manifestations of CDD (Jakimiec, Paprocka, and Śmigiel 2020; Liang et al. 2019; Wong et al. 2023; Saldaris et al. 2024). Transcriptomic analysis revealed that iNs expressed neuronal markers (MAP2, DCX) as well as a glutamatergic marker (GLUL) and did not express proliferation or progenitor makers (TOP2A, PAX6). However, the iNs did not express markers of cortical identity (FOXG1, EMX1), and expressed markers of peripheral neurons (PRPH, PHOX2B) suggesting that NGN2-induced neurons represent glutamatergic neurons but not necessarily cortical neurons, as had been previously reported (Lin et al. 2021) (Figure 1C).

**Table 1:**
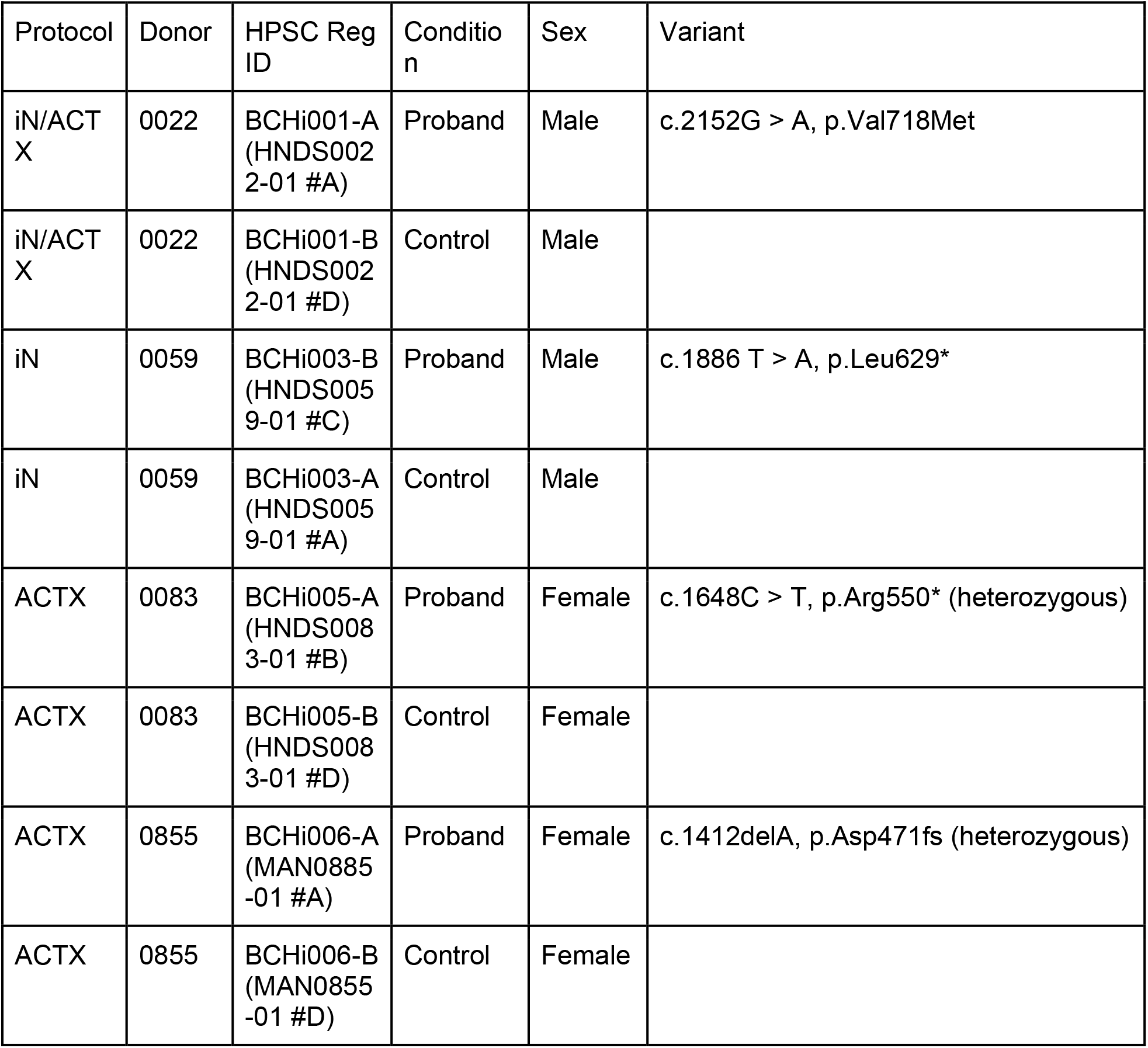
CDD Patient iPSC Lines.

| Protocol | Donor | HPSC Reg ID | Condition | Sex | Variant |
| --- | --- | --- | --- | --- | --- |
| iN/ACTX | 0022 | BCHi001-A (HNDS002 2-01 #A) | Proband | Male | c.2152G > A, p.Val718Met |
| iN/ACTX | 0022 | BCHi001-B (HNDS002 2-01 #D) | Control | Male |  |
| iN | 0059 | BCHi003-B (HNDS005 9-01 #C) | Proband | Male | c.1886 T > A, p.Leu629* |
| iN | 0059 | BCHi003-A (HNDS005 9-01 #A) | Control | Male |  |
| ACTX | 0083 | BCHi005-A (HNDS008 3-01 #B) | Proband | Female | c.1648C > T, p.Arg550* (heterozygous) |
| ACTX | 0083 | BCHi005-B (HNDS008 3-01 #D) | Control | Female |  |
| ACTX | 0855 | BCHi006-A (MAN0885 -01 #A) | Proband | Female | c.1412delA, p.Asp471fs (heterozygous) |
| ACTX | 0855 | BCHi006-B (MAN0855 -01 #D) | Control | Female |  |

**Figure 1.**
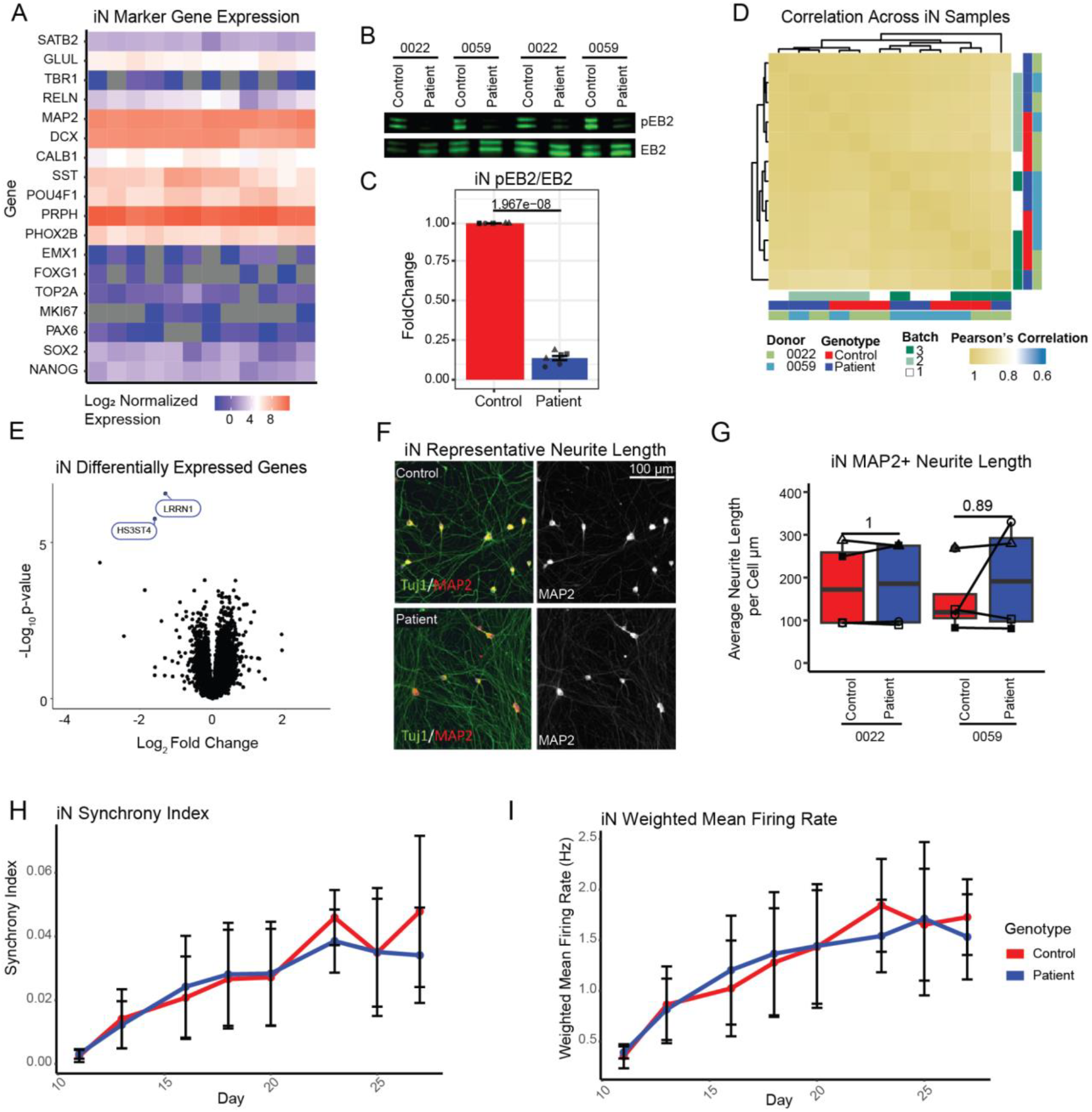
Induced Neurons Model Molecular Consequences of *CDKL5* Variants but Display NoTranscriptional, Morphological or Functional Changes. A) Normalized log fold gene expression for iN samples. B) Representative western blots for EB2 and phosphorylation of EB2 S221 with TUJ1/Beta-3-tubulin as a loading control. C) Paired t-test of phosphorylation of EB2 fold change relative to EB2 between isogenic control and CDD patient-derived neurons (N=3 for 2 donors). D) Pearson correlation coefficients for gene expression for the 5000 most variable genes between all samples from iNs. E) Volcano plot of differentially expressed genes between pooled isogenic control and CDD patient-derived samples (N=6 per genotype). F) Representative images of iN neurites immunolabeled with TUJ1 and MAP2 in both patient and isogenic control lines. A paired t-test was performed for changes in neurite length normalized to the cell body area (μm/μm^2^) for G) MAP2 immunofluorescence (N=3 batches). H) Synchrony index and I) weighted mean firing rate for INs during the first 30 days of maturation post-plating (N=4 differentiation batches, 2 donors). Error bars represent the standard error of the mean.

To confirm the loss of CDKL5 function in CDD patient neurons, we assessed phosphorylation of the known CDKL5 target, EB2 (Baltussen et al. 2018). As expected, CDD iNs had reduced levels of phosphorylated EB2 (iN, paired t-test p-value 1.97 e-8 Figure 1A). Previous human iPSC models of CDD identified an increase in mTOR signaling (Negraes et al. 2021). We also checked for changes in levels of phosphorylated AKT, phosphorylated GSK, phosphorylated S6, Acetylated tubulin and Tyrosinated tubulin, none of which had significant changes in patient derived-iNs (paired t-test p-value < 0.05) (Supplemental Fig. 1).

To understand the effects of loss of CDKL5 function on the iNs, gene expression, neuronal morphology and network activity were characterized in the male donors relative to their isogenic controls (Fig 1E-I) (Jakimiec, Paprocka, and Śmigiel 2020; Liang et al. 2019). Gene expression in CDD patient-derived iNs was strikingly similar across samples when assessed by Ampliseq, a high throughput RNA measurement platform that uses one 100 base pair sequence for counting the abundance of a transcript and is highly similar to RNAseq (Li et al. 2015).

Pearson correlations of the 5000 most variable genes were very similar across samples (R = 0.92 - 0.99) (Figure 1C). Only two differentially expressed genes (DEG) were identified consistently across lines and replicate differentiation batches: *HS3ST4* and *LRRN1*. HS3ST4 is a heparan sulfate 3-O-sulfotransferases that localizes to the presynaptic compartment and interferes with HS3ST *in vitro* inhibited hippocampal rat neuron synchrony (Maïza et al. 2020). LRRN1 is another extracellular matrix protein predicted to be involved in synapse assembly (Gene Ontology Consortium et al. 2023). Downregulation of genes associated with synaptic assembly may suggest functional differences in CDD neurons compared to control.

Since previous studies found varying deficits in neurite outgrowth, with some models showing increased outgrowth in patient-derived neurons and others revealing decreased outgrowth, we wanted to assess whether neurites in iNs were affected by loss of CDKL5 function (Negraes et al. 2021; Fuchs et al. 2022). To answer this question, iNs were imaged on the Molecular Devices IXM-C high content imaging device, and neurites were masked with the MetaXpress analysis software. In this model system, no differences were detected between patient-derived neurons and isogenic control neurons quantified by MAP2+ neurite length at 7 days post-plating (Figure 1F-H). These data suggest that the differentiation protocol and culture conditions may contribute to phenotypic differences in neurite length in these model systems.

Given the severe seizure phenotype found in CDD patients, we wanted to understand whether there were functional changes in glutamatergic neurons that could contribute to developmental and functional imbalances in patient circuits. To test this, iNs from CDD patient and control iPSC lines were plated in co-culture with healthy control astrocytes onto multi-electrode array (MEA) dishes to measure extracellular voltage changes over time.

Unexpectedly, we did not observe any measurable differences in neuronal function or network activity in the iNs with any measurement (Figure 1I-J). These negative results suggest that although iNs have some expected molecular consequences in CDD patient-derived lines, they do not display functional, morphological, or many transcriptomic alterations. There are at least two possible reasons for this negative result: 1) a potential effect of CDKL5 deficiency in neural progenitors is necessary to manifest neuronal dysfunction and 2) iN have low fidelity to cortical excitatory neurons, as previously described, and are therefore not appropriately capturing the cell types responsible for neurological manifestations (Lin et al. 2021). Modeling forebrain neurons is particularly important because CDKL5 expression is most highly enriched in these cells (Y.-C. Zhu and Xiong 2019).

To address these concerns, 3 CDD patient and isogenic control iPSC lines (2 male and 1 female) were differentiated into neural stem cells, aggregated into 3D, and differentiated into cortical neurons following an accelerated protocol that favors neurogenesis over progenitor expansion (ACTX) (Whye et al. 2023, 2024). For downstream assays, these neuron-enriched 3D cultures were dissociated and plated back into 2D for assessment of morphological and functional endpoints. Immunocytochemistry indicated that plated ACTX neurons were positive for MAP2 and TBR1 in both patient ACTX neurons and isogenic controls (Figure 2A). Further, bulk RNAseq of ACTX culture showed strong induction of cortical identity (EMX1, FOXG1) (Figure 2B). Expression of progenitor and cell cycle genes is decreased, and reliable activity in MEA suggests a protocol highly enriched for cortical neurons (Figure 2A-B, Figure 5). ACTX samples also more closely resembled intermediate progenitors and neurons from primary human fetal tissue at 14 and 18 post-conception weeks (Spearman’s R 0.32 at PCW 14 - Spearman’s R 0.29 at PCW18 for intermediate progenitors) than non-neuronal cell types (Spearman’s R 0.12 at PCW14 - Spearman’s R 0.11 at PCW 18) (Figure 2C). Though the gene expression in ACTX neurons correlate to those of interneurons and cortical neurons equally well, they express markers of early-born lower-layer neurons and Cajal-Retzius cells rather than canonical markers of inhibitory neurons, and the Spearman correlation is likely driven by pan-neuronal genes and regional markers (Figure 2C). Importantly, this is in contrast to iNs, which most closely resemble dorsal root ganglion cells (Spearman’s R = 0.2) (Figure 2C). Additionally, the ACTX neurons resemble early born cortical neurons from a previously published, reproducible human cortical organoids (Uzquiano et al. 2022) (Spearman’s R = 0.6). Characterization of ACTX neurons indicates that the updated guided protocol produces cultures of cortical glutamatergic neurons.

**Figure 2.**
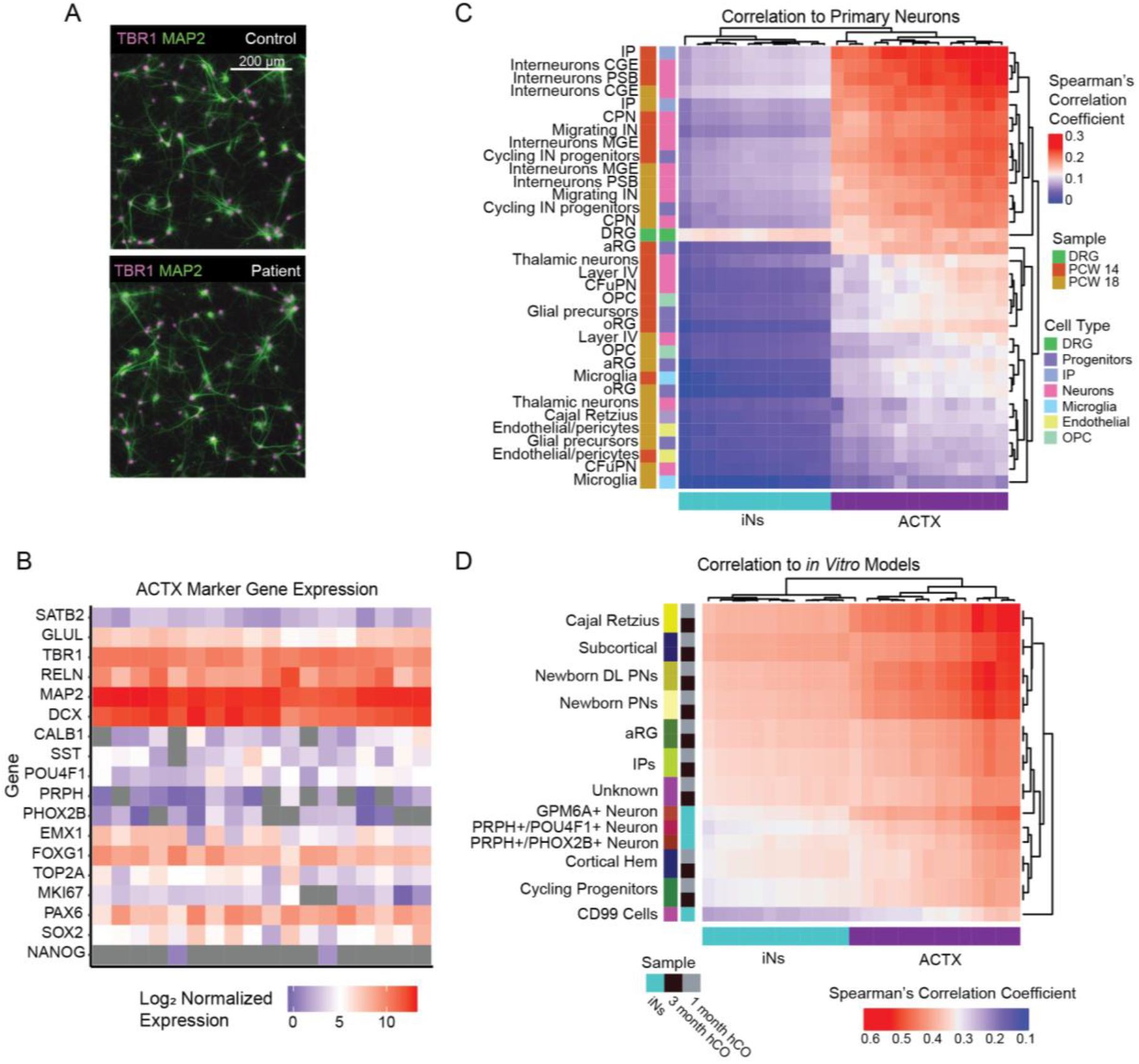
Improved Cortical Identity in ACTX Neurons Compared to iNs. A) Representative immunolabeling of TBR1+/MAP2+ ACTX neurons at day 35. B) Marker gene expression in ACTX Neurons C) Spearman correlations of the 3000 most highly expressed normalized genes counts in iNs and ACTX neurons relative to primary fetal cortex and human dorsal root ganglion. D) Spearman correlations of the 3000 most highly expressed normalized gene counts in iNs and ACTX neurons relative to published human cortical organoids at one and three months of differentiation (Uzquiano et al. 2022) and published iNs (Lin et al. 2021). Abbreviations: aRG (apical radial glia), oRG (out radial glia), IN progenitors (inhibitory neuron progenitors), CGE (caudal ganglionic eminence), MGE (medial ganglionic eminence), PSB (pallial-subpallial boundary), CFuPN (corticofugal projection neuron), CPN (cortical projection neuron), DL (deep layer), PN (projection neuron).

We next wanted to test if CDD related cellular and molecular phenotypes could be modeled in these *in vitro* cortical neuron cultures. Importantly, the ACTX neurons also capture the reduced phosphorylation of CDKL5 targets (Figure 3A-B) as demonstrated by reduced phosphorylated EB2 ratio in dissociated neurons (paired t-test p-value= 0.0014), confirming that CDKL5 function is reduced. Similar to the iNs, replicate differentiations, donors and genotypes were all very similar to each other with high Pearson’s correlation coefficients for the 5000 most highly expressed genes between samples (Pearson’s R 0.99 - 0.9) (Figure 3C). Seven DEGs were identified when pooling donors to compare CDD patients and controls (Figure 3D).

**Figure 3.**
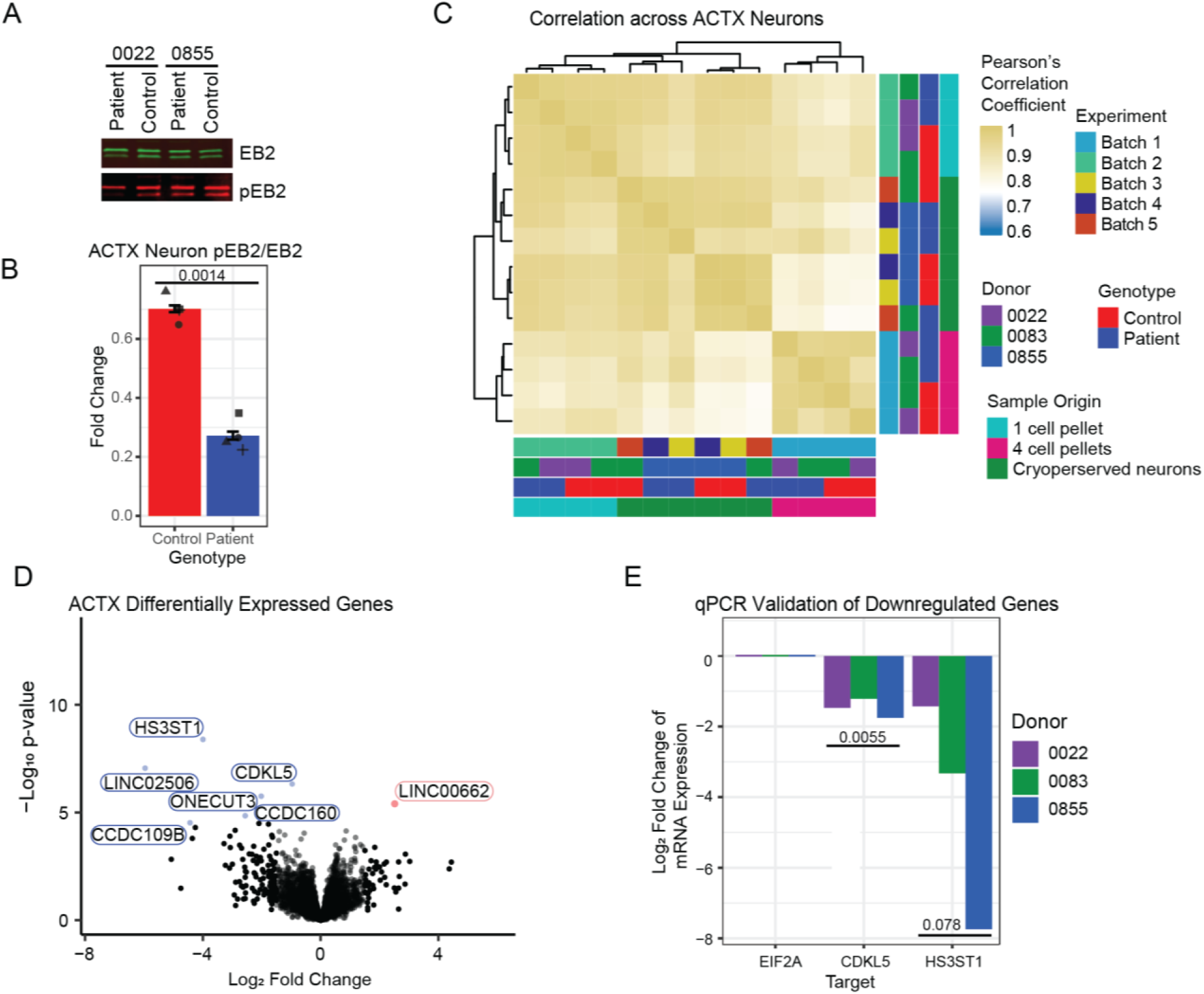
Cortical ACTX Neurons with *CDKL5* Variants Have Reduced pEB2 and Decreased Expression of *CDKL5*. A) Representative western blot of EB2 and pEB2 from ACTX neurons. B) Fold change quantification of western blots for phosphorylation of EB2 relative to EB2 across 3 donors (N = 4 donors). C) Pearson correlations across ACTX neuronal differentiations. D) Differentially expressed genes between pooled isogenic controls and CDD proband-derived neurons (N=7 sample per genotype across 3 donors). E) log2 fold change of *CDKL5* and *HS3ST1* in ACTX neurons of patient cell lines relative to isogenic controls (N=3, paired t-test). *EIF2A* is the house-keeping gene.

Interestingly, *CDKL5* had lower expression in CDD variant lines suggesting that the variants expressed in these patients reduce transcript stability. *CDKL5* pathogenic variants in these lines lead to truncations (MAN, 83, 59) and also have a known impact on splicing (22) (Olson et al. 2021; Hector et al. 2017). It has been hypothesized that CDD-associated variants causing truncation may lead to nonsense-mediated decay (Hector et al. 2017), which could explain the reduction in CDKL5 expression. Another gene that was significantly downregulated across CDD patient lines compared to controls was *HS3ST1* (Figure 3D). Other members of this gene family are involved in altering synaptic adhesion proteins and changing synaptic composition by altering the charge and affinity for protein-protein interactions in the extracellular matrix (Maïza et al. 2020), but *HS3ST1* is known as an Alzheimer’s risk gene and is involved in the cell internalization of tau (Zhao et al. 2020). HS3ST1 may be remodeling synaptic organization in ACTX neurons to contribute to hyperexcitability. Downregulation of *CDKL5* and *HS3ST1* were confirmed using qPCR on one differentiation from each donor, though only downregulation of *CDKL5* was significant (paired t-test: CDLK5 p-value= 0.0055, HS3ST1 p-value= 0.078) (Figure 3E).

Studies in mouse neurons identified microtubule-associated proteins as targets of CDKL5, suggesting that reduced CDKL5 activity and expression may lead to alteration in neurite length (Muñoz et al. 2018; Baltussen et al. 2018). Previous studies of CDD patient-derived iPSC cortical neurons with several kinds of single nucleotide variants showed increased neurite length (Negraes et al. 2021). While we did not detect differences in neurite length in the iNs, we hypothesized a similar increase in neurite length for our cohort of CDD patients using the ACTX differentiation protocol. Following dissociation from hCOs, neurons were plated into adherent culture and imaged every 6 hours for 24 hours to assess neurite length longitudinally. However, no consistent changes were found between patient-derived neurons and isogenic-controls for any donor at any time point (Figure 4) (paired t-tests: p-value > 0.05). These data suggest that CDD-associated truncation variants do not lead to changes in neurite length in lower-layer cortical neurons using these detection methods. It is possible that neurons need to be cultured in 2D for several weeks before changes in neurite length occur, as reported in previous studies (Negraes et al. 2021).

**Figure 4.**
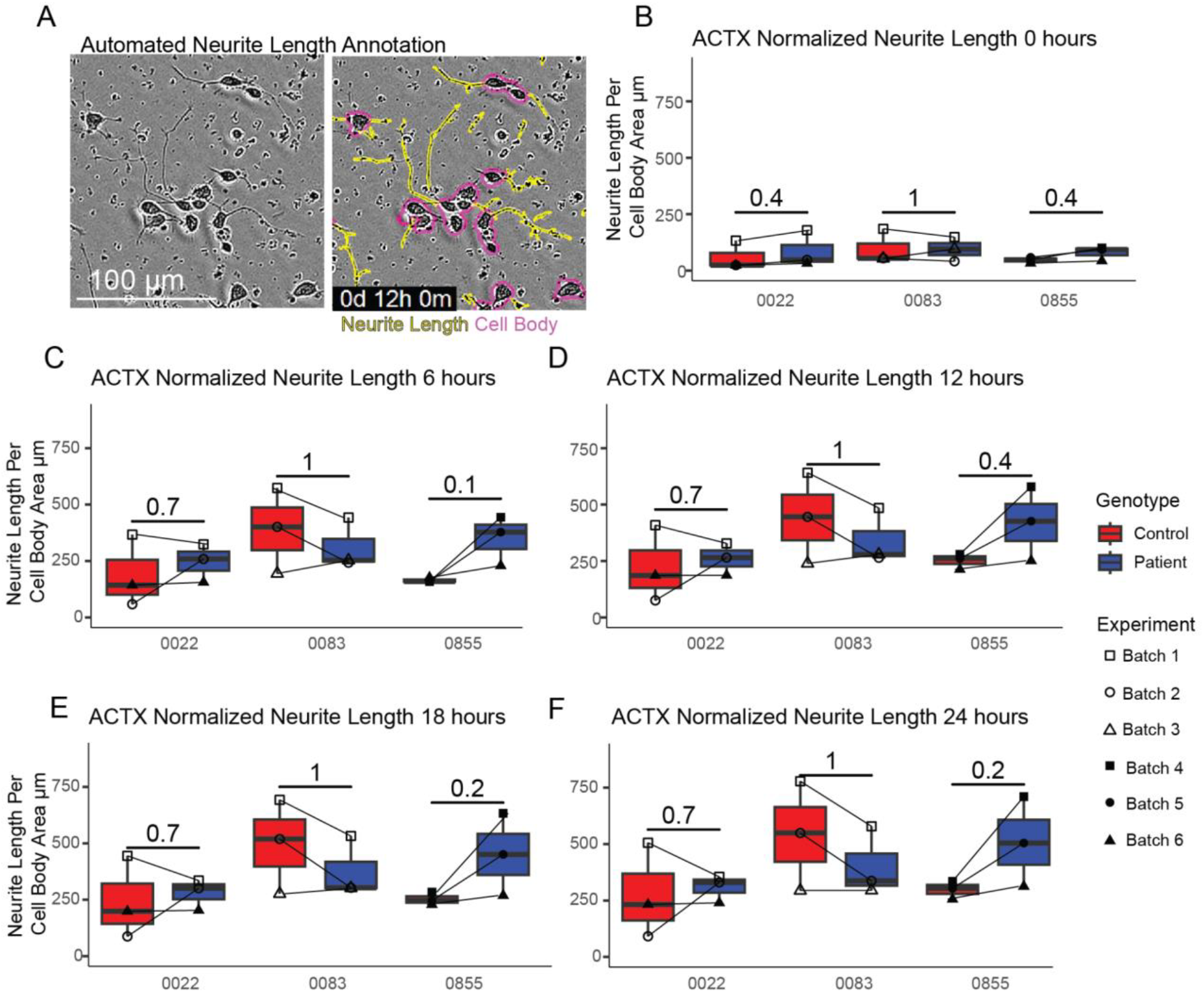
No Differences in Neurite Length between CDD Patient-derived and Control-derived ACTX Neurons. A) Example neurite and cell body segmentation at 12 hours post-plating. Quantification of normalized neurite length for each donor proband-isogenic control pair at B) 0 hours C) 6 hours D) 12 hours E) 18 hours and F) 24 hours post-plating. Post-hoc testing p-values for each donor pair are shown (N = 3).

While we did not observe functional deficits in iNs, previous studies reported cortical glutamatergic neuron hyperexcitability in 12 week old human cortical organoids and in neurons from a 2D guided differentiation (Negraes et al. 2021). Increased spontaneous excitatory postsynaptic currents and lower rheobase, indicated they are easier to excite (Negraes et al. 2021; Wu et al. 2022). We hypothesized that similar hyperexcitability would be identifiable in the ACTX neurons using MEA. By day 17, patient-derived neuron ACTX neurons had increased Synchrony Index, a measure of network connectivity, relative to controls, which persisted through day 28 (paired t-test p-value < 0.05) (Figure 5 A-C, E). Weighted mean firing rate (paired t-test p-value < 0.05) (Figure 5 D, F) and the number of spikes per burst (paired t-test p-value < 0.05) (Figure 5G) were also higher than isogenic controls, suggesting that maturing patient-derived neuron ACTX neurons are hyperexcitable and form synchronous networks sooner. MEA recordings were taken through day 70, and after 30 days of MEA recordings, no significant differences between genotypes were detected (data not shown). However, increased well-to-well variability, possibly due to neuronal clumping, cell death, or lifting, may explain this loss of effect size over time.

**Figure 5.**
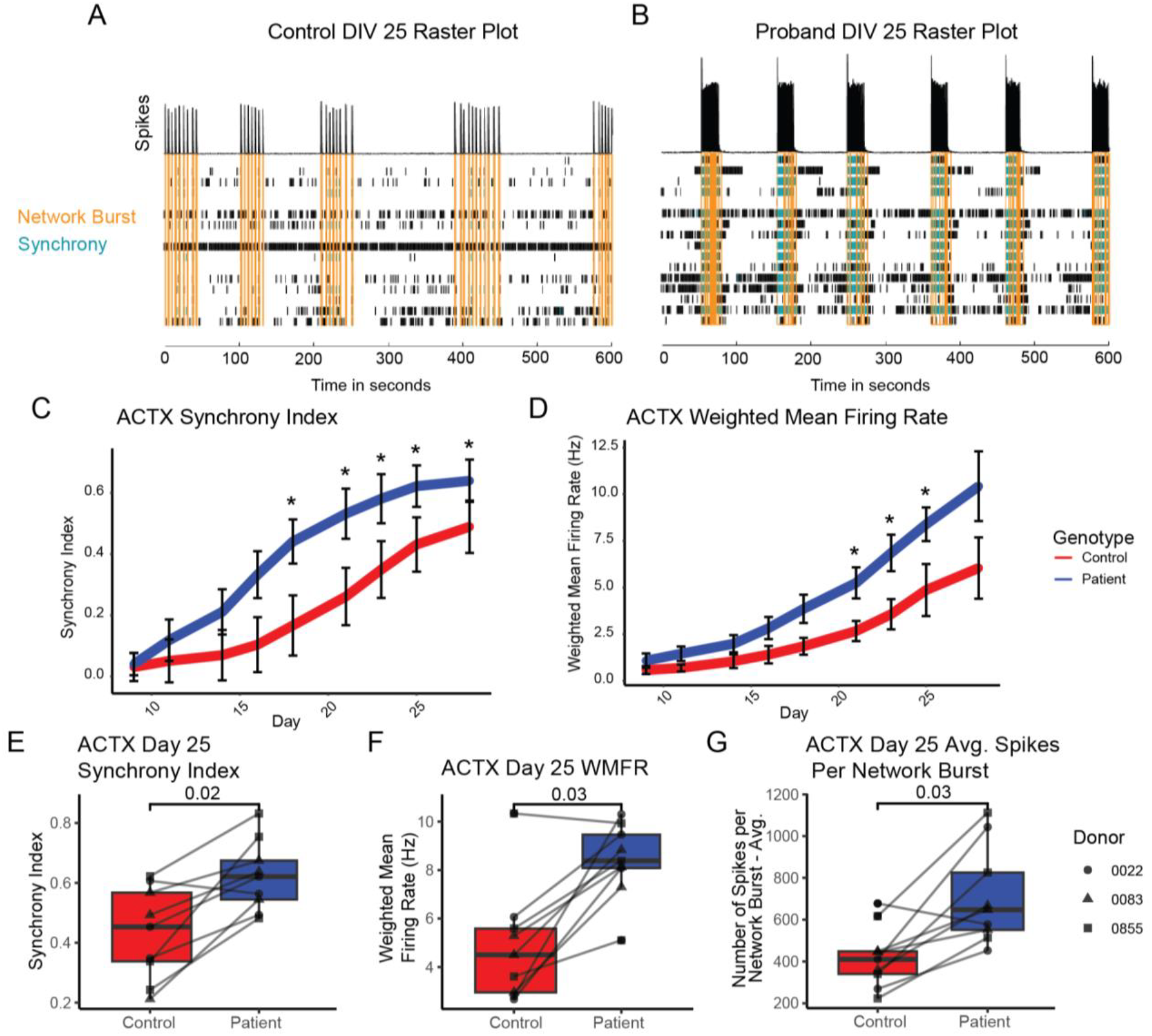
MEA Analysis Shows Early Hyperexcitability in CDD Patient-derived Neurons Compared to Control Neurons. Representative raster plots in one well at 25 days *in vitro* following hCO dissociation show network burst and synchrony in A) an isogenic control and B) a CDD proband line. C) Synchrony index for the first 30 days of recording across all donors and batches (N = 3 differentiation batches per donor across 3 donors). D) Weighted mean firing rate for the first 30 days of recording post-plating across all donors and batches (N= 3 differentiation batches per donor across 3 donors). * is p < 0.05 for paired t-test pooling across all isogenic control and proband ACTX neurons. Error bars represent the standard error of the mean. Paired t-test across pooled isogenic cases and controls at day 25 for E) Synchrony F) weighted mean firing rate (WMFR) and G) Average Spikes Per Network Burst.

## Discussion

To study cell autonomous effects of loss of CDKL5 function on glutamatergic neurons, we differentiated two CDD patient-derived iPSC lines into iNs (Zhang et al. 2013), and although these cells had phosphorylation deficits in a known CDKL5 target, no functional changes in neurons were identified. However, when 3 patient-derived iPSC lines were differentiated into cortical neurons through a guided differentiation protocol (Whye et al. 2023), CDD neurons showed an early hyperexcitability phenotype and some changes in gene expression.

Interestingly, both glutamatergic neuron models had downregulation of heparan sulfate 3-O-sulfotransferases, which are implicated in spine formation and stability (Maïza et al. 2020), supporting the previous findings that increased synapse formation and stability are found in CDD neurons (Ricciardi et al. 2012; Y.-C. Zhu et al. 2013). No changes in neurite length were detected, and these are likely independent of neuronal hyperexcitability. Importantly, our study suggests that high-fidelity human cell line models of CDD consistently identify hyperexcitability as a key pathology resulting from loss of CDKL5 function, which may be useful in future studies for development of CDD therapeutics.

We hypothesize that regional identity is the key reason for a lack of CDD-related changes in neuronal function in iNs and that a more recent direct induction protocol that includes dual SMAD inhibition or induced expression of regional transcription factors like *EMX1* to specify cortical regional identity (Nehme et al. 2018; Hulme et al. 2022; Ang et al. 2024) may have provided a better model of CDD. Our iNs, though a homogenous population, had a lower weighted mean firing rate overall (0.38 Hz for iNs; 15 Hz for ACTX at day 28), in addition to limited cortical identity. Changes in progenitor cells may not be required to identify changes in neuronal function. For example, a previous study using a CDD mouse model observed behavioral and circuit disruptions when global CDKL5 deficiency was induced postnatally, and postnatal genetic restoration of CDKL5 was also sufficient to ameliorate behavioral changes (Terzic et al. 2021). Their results suggest that CDKL5 deficiency in neurons is sufficient to elicit at least some cellular changes. We encourage future studies to rigorously define their *in vitro* cell types prior to interpretation.

The observation that there were no consistent changes in neurite length when comparing patient and control iNs and ACTX neurons was unexpected given that one of the known targets of CDKL5 kinase function is the microtubule associated protein, MAP1S (Baltussen et al. 2018). However, it is possible that specific maturation states or culture conditions are needed to quantify these deficits. Previously reported *in vitro* findings that neurite length varied between patient and control derived neurons were derived from neurons cultured for 8 weeks and may be dependent on extended maturation times or an upper-layer neuron population (Negraes et al. 2021). Postmortem analysis of neurite length from CDD patients also showed inconsistencies in neurite length differences across donors (Negraes et al. 2021). It is possible that alterations in neurite length are specific to variants between the ATP-binding site and serine-threonine kinase active site (Negraes et al. 2021) and not the truncating variants included in this study, or could be a consequence of differences in the model systems. If the differences in *CDKL5* variants are responsible, this data provides evidence for some genotype-phenotype correlations in CDD patients (Olson et al. 2019; MacKay et al. 2021; Bahi-Buisson et al. 2012).

Neuronal hyperexcitability has been identified as a consistent feature in mouse models of CDD. There are several proposed mechanisms leading to hyperexcitability, including changes in phosphorylation of CaV2.3 leading to slower inactivation (Sampedro-Castañeda et al. 2023), as well as an upregulation of AMPA receptors (Yennawar, White, and Jensen 2019) and NMDA receptors (Okuda et al. 2017; Tang et al. 2019) at the synapse. Since there have been challenges reproducing spontaneous seizure phenotypes in young mice lacking *Cdkl5* expression, researching neuronal hyperexcitability mechanisms in human model systems is of particular interest. A first look into increased intrinsic excitability in human iPSC-derived cortical neurons found an increase in K+ and Na+ current density (Wu et al. 2022). Our study shows that hyperexcitability in CDD is consistent in human iPSC-derived cortical neuron models. Future work identifying the molecular mechanisms leading to hyperexcitability in human model systems, and potential ways to rescue neuronal function, are an exciting prospect for CDD research.

## Figures

**Supplemental Figure 1.**
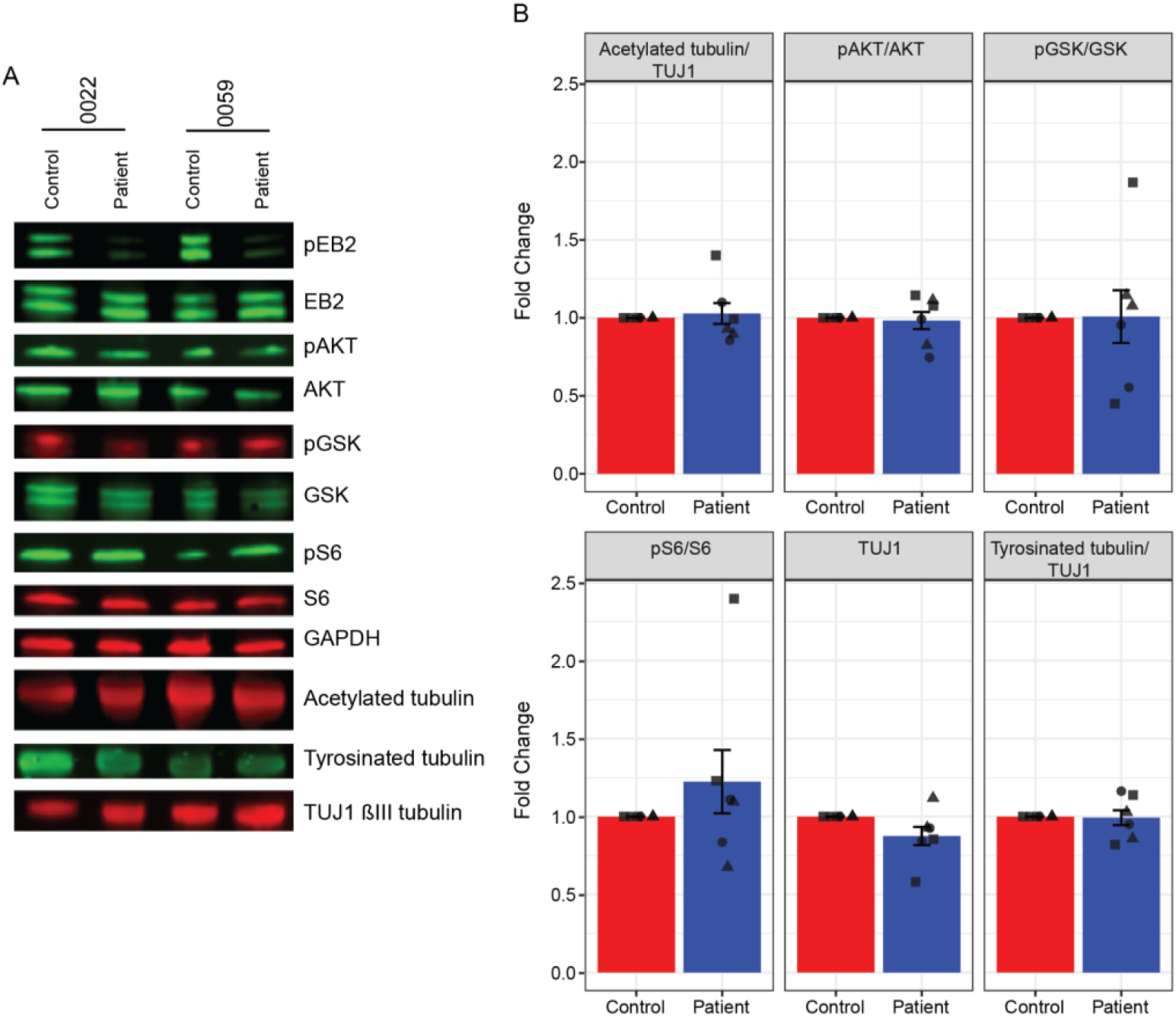
No Altered mTOR Signaling in iNs. A) Representative Western blots for phosphorylation of different mTOR-signaling associated proteins. pEB2 and EB2 as shown in Figure 1 for reference. B) Quantification of normalized protein abundance for each tested protein (N=3). Each shape represents a differentiation batch.

## Methods

### Data Availability

Scripts used for transcriptomic analysis and plotting will be available on Github following acceptance. Transcriptomic data will be deposited on dbGaP following acceptance.

### iPSC maintenance, quality control

All iPSC lines were cultured in Stemflex (Thermofisher #A3349401) or in mTeSR Plus (StemCell Technologies #100-0276) and Matrigel (Corning Inc #354277) or Geltrex (Thermofisher Scientific #A1413301) coated plates. iPSC lines had been previously karyotyped and no abnormalities were detected. All iPSCs were used for neuronal differentiation within 10 passages of the previously published normal karyotypes (Chen et al. 2021). All cell cultures were routinely tested for mycoplasma using luminescence (MycoAlert Lonza #LT07-318).

### iN differentiation

All iNs were cultured in mTeSR. NGN2-induction was performed as previously described (Zhang et al. 2013). iPSCs were dissociated in Accutase (Innovative Cell Technology #AT 104– 500) and seeded on Geltrex-coated plates (Thermo Fisher Scientific #A1413301). The next day, the media was changed to N2 Medium and supplemented with DOX (2 *μ*g/mL, Millipore #324385–1GM), BDNF (10ng/mL, Thermo Fisher Scientific #17502-048), NT3 (10 ng/mL, Peprotech #450–03), Laminin (5 *μ*g/mL Life Technologies #23017-015). The following day cultures underwent puromycin (1 *μ*g/mL, Invitrogen #ant-pr-1) selection for 2 days. Cells were switched to B27 Medium on day 2, supplemented with Ara-C (2 *μ*M, Sigma-Aldrich #C1768), and DOX, BDNF, NT3 and Laminin as previously described until day 6. After day 6, iNs were co-cultured with 11,250 human astrocytes (Ncardia, K0101 Ncyte Astrocyte Kit II) or in astrocyte conditioned media.

### ACTX differentiation

Differentiation of adherent neural stem cells followed by 3D aggregation for neuron differentiation is adapted from a previously published protocol (Whye et al. 2024). For 2D neural induction, iPSCs were grown to near confluence (90-95%) in either Stemflex (Thermofisher #A3349401) or in mTeSR Plus (StemCell Technologies #100-0276) prior to dissociation into single cells using Accutase (Innovative Cell Technologies #AT 104–500). iPSCs were seeded into either Matrigel-coated (Corning Inc #354277) or Geltrex-coated (Thermofisher Scientific #A1413301) tissue culture-treated vessels at a density of 150K cells/cm^2^. Enhanced neural induction was accomplished using dual Smad inhibitors, 10 *μ*M SB431542 (Selleck Chemicals #S1067) and 0.25 *μ*M LND193189 (Tocris Bioscience #TB6053-GMP), along with FGF receptor inhibitor, 5*μ*M SU-5402 (Selleck Chemicals #S7667) and WNT inhibitor, 5 *μ*M XAV939 (Selleck Chemicals #S1180). After 7 days, adherent neural stem cells were dissociated into single cells using Accumax (Innovative Cell Technologies #AT 104–500) and seeded in 1x CEPT chemical cocktail (Tocris Bioscience #7991) along with the morphogens epidermal growth factor, 20 ng/mL EGF (Peprotech #AF-100-15) and basic fibroblast growth factor, 20 ng/mL bFGF or FGF-2 (Peprotech #AF-100-18B) at 125,000 cells/cm^2^. Neural stem cell culture was expanded for one or two weeks. To generate suspended 3D cultures, singularized neural stem cells were seeded at a density of 5,000,000 cells/well in 6-well, tissue culture plates (Corning Inc.) that were coated with the anti-adhesive solution Pluronic F-127 (Sigma-Aldrich # P2443-250G), and plates were placed on an LSE Orbital Shaker (Corning) at 100 rpm. Aggregated neural spheroids continued to receive expansion medium until day 21, at which point their medium was replaced with differentiation medium containing the Wnt-activating compound, 3 *μ*M CHIR99021 (Tocris Bioscience # 4423) and the MEK inhibitor, 2.5 *μ*MPD0325901 (Selleck Chemicals #S1036) for 7 days. At D28, differentiation medium was replaced with terminal acceleration medium containing Notch/gamma-secretase inhibitors DAPT 10 *μ*M (Selleck Chemicals #2634), Compound E 100nM (Tocris #6476), and 1x Culture One supplement (Thermofisher #A33202-01) along with the neurotrophic factors 20 ng/mL BDNF (Peprotech #450-02), 20 ng/mL NT-3 (Peprotech #78074.1), and 20 ng/mL NGF (Peprotech #256-GF-100/CF). End-stage cortical organoids were dissociated with Accumax and plated as single cells at 125K cells/cm^2^ in vessels for immunofluorescence staining, living imaging or multielectrode array analysis. Dissociated neurons were cultured in BrainPhys (StemCell Technologies #05790) supplemented with BDNF, NT3, NGF as described above and Laminin (Life Technologies, cat. no. 23017-015).

### Western Blots

Cell cultures were lysed in RIPA buffer with 1:100 PMSF (ChemCruz, 200 mM), phosphatase inhibitor (Selleckchem, 100 mM), and proteinase inhibitor (ChemCruz). Undissolved materials were removed using benchtop microcentrifuge (4 °C, 14,000 rpm, 20 min). Sample protein contents were measured by Pierce™ BCA Protein Assay Kit (catalog #23225; Thermo Scientific). Equal amounts of protein were loaded and resolved by polyacrylamide gel electrophoresis (PAGE) using Criterion TGX 4–20% Precast Gels (Biorad) and transferred onto PVDF membrane using iBlot gel transfer system (Invitrogen). Membranes were dried overnight and rehydrated with 100% Methanol, MilliQ Water, and TBS before blocking for one hour at room temperature in Intercept (TBS) blocking buffer (LI-COR). Membranes were then incubated for one hour at room temperature with EB2 1:500 (Abcam ab234843), 1:500 p-EB2 S222 (CovalAb pab01032-P) Signaling Technologies #5364), 1:500 S6 (Santa Cruz #sc-74459), 1:500 p-AKT (Sigma #05-1003), 1:500 AKT (Cell Signaling Technologies #4691S), 1:500 p-GSK (Cell Signaling Technologies #5558), 1:500 GSK (Cell Signaling Technologies #5676S) 1:4000 Acetylated tubulin (Sigma #T7451), 1:4000 TUJ1 (Sigma #T8660), 1:4000 Tyrosinated tubulin (Millipore #MAB 1864), 1:2000 GAPDH (Invitrogen #AM4300), washed by TBS-T, incubated for one hour at room temperature with secondary antibodies at 1:10000 (Alexa Fluor488, ThermoFisher #A28175; Alexa Fluor568, ThermoFisher #A-11011) and imaged using the Odyssey imaging system (LI-COR). Images were processed and quantified using Image Studio acquisition software (LI-COR). Protein fluorescent intensity was normalized to a total protein stain (Licor 926-11011) as an internal control and then compared between proband and isogenic controls. 3 unique differentiation batches were used for each proband-isogenic control pair for both iNs and ACTX neurons. Unprocessed western blots are attached as a supplementary file.

### Ampliseq Library Preparation, Preprocessing and Analysis

Total RNA was harvested from iNs 3 weeks post-induction (Takara Bio catalog #740971) and converted into cDNA using SuperScript IV VILO (Thermofisher Scientific #11756050). The Ion Ampliseq Transcriptome Human Gene Expression Kit (Thermo Fisher Scientific #A31446) in an automated Ion Chef instrument was used to generate libraries that were then sequenced on an IonTorrent S5 Sequencer (Thermofisher Scientific). The Torrent Suite was used to preprocess the data and generate a matrix of read counts for each sample and gene probe. Ampliseq read counts were normalized to counts per million. Differential gene expression analysis was performed using Limma’s default settings (Ritchie et al. 2015). The union of the 5000 most highly expressed genes from both samples was used to calculate Pearson’s correlation coefficients across all samples with normalized read counts.

### Bulk RNAseq Library Preparation, Preprocessing and Analysis

RNA was extracted from day 35 neurons using (Takara Bio catalog #740971) from either frozen cell pellets or cryopreserved neurons. Libraries were prepared from 125 ng of total RNA using the NEGEDIA Digital mRNA-seq research grade sequencing service v2.0 (NEGEDIA Srl.) (Xiong et al. 2017), which included library preparation, quality assessment, and sequencing on a NovaSeq 6000 sequencing system using a single-end, 100 cycle strategy (Illumina Inc.) The raw data were analyzed by NEGEDIA Srl. proprietary NEGEDIA Digital mRNA-seq pipeline (v2.0) which involves a cleaning step by quality filtering and trimming, alignment to the reference genome and counting by gene (Xiong et al. 2017; Bushnell 2014; Anders, Pyl, and Huber 2015). The raw expression data were normalized (v1.2.0) (Subramanian et al. 2005). RNA extracted directly from neurons cryopreserved after 11 months in cold storage generated higher duplicate reads and was removed from further analysis. Bulk RNAseq was analyzed using DeSEQ2 (Love, Huber, and Anders 2014) to identify differentially expressed genes between CDD variant lines and controls. Raw read counts were used as input, and genes with fewer than 10 reads were removed from the analysis. The union of the 5000 most highly expressed genes from both samples was used to calculate Pearson’s correlation coefficients across all samples with normalized read counts.

### Spearman Correlations to External Datasets

All referenced external datasets are from public single-cell RNAseq atlases and were analyzed for this study using Seurat (v5.1) (Hao et al. 2024). For the primary human fetal tissue, read counts were aggregated across cell types (Seurat Aggregate Expression Function v5.1) that had less than 10% mitochondrial reads in each cell and organoid samples were filtered to include only cells from cell lines without known pathogenic variants (Single Cell Portal ID: SCP1756) (Uzquiano et al. 2022; Tarhan et al. 2023). All library preparations were pooled together in the human dorsal root ganglion data prior to aggregation (Nguyen et al. 2021) (GEO ID: GSE168243). External datasets are SCT transformed prior to aggregation (Choudhary and Satija 2022). Normalized read counts were used for iN or ACTX transcriptomic data. Spearman correlations were performed with 3000 highest expressed genes.

### qPCR

RNA was extracted from frozen cell pellets (Takara Bio catalog #740971) and converted into cDNA (LunaScript RT Supermix Kit New England Biolabs E3010S). Approximately 500 ng of cDNA was used per reaction (Luna Universal qPCR Master Mix New England Biolabs: M3003X; Microamp EnduraPlate Thermo Scientific: 448354) performed using a QuantStudio 3 (Thermo Fisher Scientific). *EIF2A* was used as the housekeeping gene. Primers: *HS3ST1*_fwd: 5’ AACTGGCTGCGCTTTTTCC 3’, *HS3ST1*_rev: 5’ ACCTCTCGACCTTTTGGATCT 3’, *CDKL5*_fwd: 5’ ATTACACAGAGTACGTTGCCAC 3’, *CDKL5*_rev: 5’ CCACGGACTTTCCATAGGGAG 3’, *EIF2A*_fw: 5’ TGGTGTCATCGAGAGCAACTG, *EIF2A*_rev: 5’ GGCTTCTCAAAACCGTAAGCA 3’. Primer bank IDs: 52426776c3, 83367066c3, 126091140c2, respectively.

### Immunocytochemistry for Neurite Length

The fixed and stained plates with cells were imaged with ImageXpress MicroXLS Widefield High-Content Molecular Device Microscope (Molecular Device, LLC, MetaXpress v.6.6.2.46). Cells were fixed in 4% paraformaldehyde for 10 minutes at room temperature. Nonspecific staining was blocked with 5% NGS, 2% BSA, 0.1% Triton X-100 in dPBS for one hour at room temperature. Primary antibodies were diluted in 5% NGS, 2% BSA, 0.1% Triton X-100 in dPBS and incubated on the plates overnight at +4 °C. 1:2000 MAP2 (Abcam #5392) and 1:500 TBR1 (Abcam # 31940). The next day, the cells were washed with dPBS and incubated with secondary antibodies for one hour at room temperature. Secondary antibodies were diluted 1:500 (Alexa Fluor488 ThermoFisher #A28175 and Alexa Fluor568 ThermoFisher #A-11011). Nuclei were stained with Hoechst 33258 (Thermo Fisher Scientific # H3569) at 1:500 dilution together with secondary antibodies.

### Live Imaging

Dissociated neurons were plated at 40,000 cells per well of a 96-well plate in terminal acceleration media. All neurons were imaged and analyzed using the IncuCyte S3 Live cell analysis imaging system and associated software (Sartorius V2018B) and imaged every 6 hours for the first 24 hours following plating. Phase images were acquired with the IncuCyte® ZOOM 20X Objective (#4464).

### Neurite Length

Following imaging, analysis was completed using the Incucyte® Neurotrack Analysis Software Module. The algorithm masks and quantifies each image and returns neuronal cell metrics, including neurite length, cell body area, and number of cell clusters directly on phase contrast images. Measurements were exported and plotted using custom R scripts.

### MEA Plating and Analysis

On Day 6, 75,000 iNs were replaced with 11,250 human astrocytes (Ncardia, K0101 Ncyte Astrocyte Kit II) following manufacturer’s instructions onto 48 well or 6 well low-density (LD)-MEA platform plates (Axion BioSystems) in B-27 media. 10 *μ*M Y27632 (Cayman Chemical Company #10005583) was supplemented into the media for one day to improve cell survival. Recordings began on day 8 post-plating and cultures were recorded every 2-3 days on the MaestroPro multi-electrode array system (Axion Biosystems). The number of active electrodes per well did not differ between cases and controls, and wells with fewer than 10 active electrodes by day 11 were excluded.

Following dissociation from hCOs, 100,000 ACTX neurons per well were seeded on 48 well LD-MEA plates coated with 0.10% PEI (Sigma-Aldrich, #408727-250g) and 5 - 100 *μ*g/mL of laminin (Life Technologies,#23017-015). ACTX neurons were co-cultured with 12,000 - 15,000 human astrocytes (Ncardia, K0101 Ncyte Astrocyte Kit II).

All activity was spontaneous and recorded for 10 minutes per plate. To export the data, recordings were batch-processed, and .spk files were then transferred into the Neural Metrics tool (Axion BioSystems) for further analysis. All metrics for all days were exported as .csv files and imported into R for analysis. Wells with fewer than 10 active electrodes by day 11 were excluded.

## Acknowledgements

We would like to thank Ivy Pin-Fang Chen, Cidi Chen, and all members of the Human Neuron Core at Boston Children’s Hospital (P50HD105351). We are grateful to Drs. Maria Sundberg, Kellen Winden, and Wardiya Afshar-Saber, and members of the Sahin lab for critical reviewing of the manuscript and helpful comments. Finally, we would like to thank the patients and families who contributed to this work.

Madison R. Glass: Writing-original draft, Visualization, Formal analysis. Dosh Whye: Conceptualization, Investigation, Methodology, Data curation, Writing - review & editing. Nickesha Anderson: Conceptualization, Investigation, Data curation. Delaney Wood: Investigation, Data curation. Nina R. Makhortova: Investigation, Data curation, Writing - review & editing. Taryn Polanco: Investigation, Data curation, Writing - review & editing. Kristina H. Kim: Investigation, Data curation. Kathleen E. Donovan: Investigation, Data curation. Lorenzo Vaccaro: Investigation, Data curation, Formal Analysis. Ashish Jain: Formal Analysis. Davide Cacchiarelli: Investigation, Data curation. Liang Sun: Formal Analysis. Elizabeth D. Buttermore: Conceptualization, Investigation, Data curation, Writing - original draft, Writing - review & editing, Funding acquisition, Supervision, Project administration. Mustafa Sahin: Conceptualization, Investigation, Data curation, Writing - original draft, Writing - review & editing, Funding acquisition, Supervision, Project administration.

